# Opposing Autonomic Pathways Link Childhood Trauma to Mental Imagery Deficits

**DOI:** 10.64898/2026.01.10.698789

**Authors:** Juha Silvanto, Wenyue Gao, Yoko Nagai

**Affiliations:** Center for Cognitive and Brain Sciences, University of Macau, China SAR; Department of Psychology, Faculty of Health Sciences, University of Macau, China SAR; Brighton & Sussex Medical School, Department of Neuroscience, University of Sussex, UK

## Abstract

Although childhood trauma alters neurobiological systems supporting emotion regulation and autonomic function, mechanisms linking trauma to mental imagery deficits remain unexplored. Here we examined whether autonomic nervous system symptoms mediate trauma-imagery associations in a general population sample (n = 433). All trauma types showed associations with reduced imagery vividness (r = -.12 to -.23), with emotional neglect and emotional abuse demonstrating the strongest effects. Physical trauma types and emotional abuse correlated with increased autonomic symptoms (r = .35-.38). Parallel mediation analyses revealed that autonomic symptom burden (BPQ ANS subscale), but not anxiety, significantly mediated trauma-imagery relationships through opposing mechanisms. Physical trauma types and emotional abuse showed negative indirect effects through heightened autonomic reactivity, with complete mediation for emotional abuse and physical neglect. In contrast, emotional neglect showed a positive indirect effect through reduced autonomic reactivity (a pattern that emerged when controlling for other trauma types) alongside the strongest direct effect on imagery. These findings demonstrate that childhood trauma impacts mental imagery through distinct autonomic mechanisms—hyperreactive dysregulation for physical/emotional abuse versus hyporeactive blunting for emotional neglect—suggesting autonomic-targeted interventions should be tailored to trauma type.

## Introduction

Childhood trauma affects up to 50% of the population worldwide and represents one of the most significant risk factors for lifelong psychological and physical health problems (Felitti et al., 1998; Hughes et al., 2017). Adverse childhood experiences encompass a broad range of maltreatment including emotional, physical, and sexual abuse, as well as emotional and physical neglect, household dysfunction, and exposure to violence. These experiences fundamentally disrupt the developing child’s capacity to form coherent internal representations of self, others, and the world, occurring during critical periods when neural systems responsible for stress regulation, attachment, and self-awareness are rapidly developing (van der Kolk, 2014).

Early adversity produces lasting alterations in neurobiological systems including the brain circuits supporting emotion regulation and cognitive function (Teicher & Samson, 2016; McLaughlin et al., 2014). Childhood trauma sensitizes threat-detection systems, leading to persistent hypervigilance to bodily signals experienced as threatening (Schaan et al., 2019; Dale et al., 2022). This dysregulation is particularly evident in amplified cardiovascular and respiratory responses, which contribute to chronic anxiety and trauma-related disturbances (Schmitz et al., 2011).

Among the cognitive processes vulnerable to these trauma-induced alterations, mental imagery may be particularly susceptible due to its fundamental dependence on integrated bodily and sensory representations. Mental imagery (the capacity to generate and manipulate sensory-like representations in the absence of corresponding external stimuli) plays a fundamental role in episodic memory, planning, emotional regulation, and social cognition (Kosslyn et al., 2001). Generating vivid images has been proposed to require embodied simulation of multisensory experiences, drawing on stored representations of sensory experiences, body states, and contextual information (Silvanto & Nagai, 2025; Muraki et al., 2023). In this view, interoceptive signals may serve as a physiological anchor for imagery, ensuring that internally generated experiences remain vivid and embodied rather than purely abstract.

Recent empirical evidence supports this vulnerability. Gao et al. (2025) found that individuals with acquired imagery loss (known as aphantasia) reported significantly higher childhood trauma exposure than those with intact imagery. Furthermore, the acquired aphantasia group self-reported elevated levels of autonomic symptom burden, raising the possibility that childhood trauma may impact imagery through autonomic dysregulation. According to interoceptive models, chronic autonomic dysregulation produced by trauma (Dale et al., 2022; Teicher & Samson, 2016) may interfere with the integration of bodily and sensory information needed for mental imagery. It may do so by introducing excessive interoceptive “noise” that disrupts the precise bodily signals required for imagery generation, or by promoting dissociative disconnection from bodily awareness as a defensive response to overwhelming autonomic arousal (Lanius et al., 2010), thereby removing the interoceptive scaffold that anchors vivid mental simulation.

However, the nature of this dysregulation may be more nuanced than a simple increase in autonomic reactivity. Optimal mental imagery generation may require maintaining an appropriate dynamic range of autonomic nervous system regulation; interoceptive signals must be sufficiently robust to anchor embodied simulation, yet regulated enough to support the precise neural computations required for stable imagery (Nagai et al., 2025). Evidence suggests that deviations in either direction may be detrimental: while acquired aphantasia following trauma is associated with elevated autonomic symptoms (Gao et al., 2025), congenital aphantasia (present from birth) shows the opposite pattern, i.e. reduced interoceptive sensitivity (Monzel et al., 2021). This raises a critical question for understanding trauma-related imagery deficits: do different types of childhood trauma produce distinct patterns of autonomic dysregulation, with corresponding effects on imagery through divergent pathways? Some forms of trauma might drive autonomic function toward hyperreactivity, others toward hyporeactivity, both representing departures from the optimal range necessary for vivid imagery. Identifying such trauma-specific autonomic-imagery pathways could clarify mechanisms underlying individual differences in trauma outcomes.

Despite this emerging evidence, critical questions remain. First, whether trauma-imagery associations extend beyond aphantasia to the normal range of imagery ability is unknown. Second, it remains unclear whether trauma impacts imagery specifically through autonomic dysregulation or through broader affective mechanisms such as generalized anxiety. Third, if the autonomic mechanism is valid, different trauma types should show opposing patterns of mediation (hyperreactive versus hyporeactive pathways) reflecting the proposed bidirectional dysregulation. The current study addressed these questions with three aims. First, we examined associations between specific childhood trauma types and mental imagery vividness in a general population sample. Second, we tested whether self-reported autonomic symptom burden mediates trauma-imagery relationships, and whether different trauma types show opposing patterns of mediation. Third, we assessed whether autonomic symptoms provide a more specific mediating pathway than general anxiety.

## Methods

### Participants

The sample comprised 433 adults (63.3% women, n=274; 36.5% men, n=158; 0.2% non-binary, n=1) aged 18-75 years (see Table 1 for mean and SD), recruited online via Profilic platform without restrictions or criteria on imagery ability. All participants provided informed consent, and the study was approved by the local ethics committee at University of Macau. Participants were paid 5 USD for their participation.

**Table 1.** Descriptive Statistics.

| Variable | M | SD | N |
| --- | --- | --- | --- |
| Age (years) | 51.5 | 15.3 | 431 |
| BPQ Body Awareness | 60.8 | 19.8 | 422 |
| BPQ Autonomic Scale (ANS) | 41.2 | 14.6 | 425 |
| GAD-7 | 6.70 | 5.66 | 419 |
| CTQ Emotional Abuse | 9.41 | 5.10 | 419 |
| CTQ Sexual Abuse | 7.39 | 4.70 | 419 |
| CTQ Physical Abuse | 7.85 | 4.36 | 419 |
| CTQ Emotional Neglect | 10.9 | 5.45 | 419 |
| CTQ Physical Neglect | 8.11 | 3.68 | 419 |
| PSIQ Total score | 7.13 | 2.09 | 424 |

### Materials and Procedure

Participants completed all measures online via the Qualtrics survey platform in a single session lasting approximately 20-25 minutes. Questionnaires were presented in randomized order to control for order effects. To ensure data quality, two attention check items were embedded within the survey (one in the AFS and one in the BPQ-SF). These items instructed participants to select a specific response option (e.g., “Please select ‘rarely’ for this item”) to verify attentive responding. As in our prior work (Gao et al, 2025), Participants who failed both attention checks or had response times below 2 seconds per item on any questionnaire were excluded. Specifically, the following questionnaires were used:

#### Plymouth Sensory Imagery Questionnaire (PSI-Q)

Participants completed the PSI-Q (Andrade et al., 2013), which assesses mental imagery vividness across seven modalities: visual, auditory, olfactory, gustatory, tactile, bodily sensations, and emotional imagery. This 35-item instrument asks respondents to rate their imagery experiences for each sensory domain.

#### Childhood Trauma Questionnaire (CTQ)

Adverse childhood experiences were measured using the CTQ (Bernstein et al., 1998), a 28-item retrospective assessment covering five maltreatment types: physical abuse, emotional abuse, sexual abuse, physical neglect, and emotional neglect. Items use a 5-point scale ranging from 1 (“Never true”) to 5 (“Very often true”). Given the sensitive content, participants were told they could omit questions without withdrawing from the study.

#### Body Perception Questionnaire – Short Form (BPQ-SF)

The BPQ-SF (Cabrera et al., 2018) was administered to evaluate interoceptive awareness across multiple dimensions. This measure includes subscales assessing body awareness (awareness of body processes and sensations) and autonomic nervous system symptoms (BPQ ANS; frequency of autonomic reactivity symptoms across cardiovascular, gastrointestinal, respiratory, and thermoregulatory systems). Respondents indicate how frequently they notice each symptom using a 5-point scale.

#### Generalized Anxiety Disorder Scale (GAD-7)

Anxiety symptoms were assessed using the GAD-7 (Spitzer et al., 2006), which contains 7 items measuring generalized anxiety over the preceding two weeks. Response options range from 0 (“not at all”) to 3 (“nearly every day”), yielding total scores between 0 and 21. Severity categories are: minimal (0–4), mild (5–9), moderate (10–14), and severe (15–21).

### Statistical analyses

All analyses were conducted using jamovi (version 2.3.28). Missing data were handled using listwise deletion (final sample being n= 414-419 depending on the measure). Significance was set at α = .05 (two-tailed). We first report bivariate correlations between all study variables to examine zero-order associations before testing multivariate models.

#### Hierarchical Regression

We conducted hierarchical multiple regression to further examine the link between trauma and imagery. Model 1 included only demographics (age, gender) to establish baseline variance; Model 2 added all five childhood trauma subscales (CTQ) to test whether trauma predicts imagery deficits; Model 3 added interoceptive symptoms (BPQ-Body Awareness, BPQ ANS) and current anxiety (GAD-7) to examine whether these symptoms explain additional variance beyond trauma, providing the foundation for subsequent mediation analyses.

#### Mediation Analyses

parallel mediation with both BPQ ANS subscale and GAD-7 as simultaneous mediators (trauma → BPQ ANS/GAD-7 → imagery) was carried out, with all five CTQ subscales as simultaneous predictors with PSIQ as outcome. The BPQ ANS subscale was selected as it specifically captures awareness of autonomic nervous system activity, theoretically most relevant to trauma-related interoceptive dysregulation. GAD-7 was included to distinguish general anxiety symptoms from somatic-interoceptive dysregulation, testing whether trauma operates through cognitive worry versus bodily awareness pathways. Indirect effects were tested using bias-corrected bootstrap confidence intervals (10,000 resamples), considered significant when 95% CIs excluded zero. We report unstandardized coefficients (B) with standard errors and standardized coefficients (β) for effect size comparison.

## Results

Descriptive statistics for all measures are shown in Table 1. Table 2 presents bivariate correlations among all key study variables. Higher trauma exposure was associated with reduced imagery vividness across all trauma types, with the strongest effects observed for emotional neglect (r = -.23, p < .001) and emotional abuse (r = -.21, p < .001). Imagery was also significantly negatively correlated with anxiety (GAD-7; r = -.25, p < .001) and with BP_ANS (r = -.24, p < .001). GAD was significantly associated with all CTQ subscales, with correlations ranging from r = .20 (emotional neglect) to r = .37 (emotional abuse). BP_ANS showed similar significant associations with all trauma types except emotional neglect. Childhood trauma subscales showed moderate-to-large intercorrelations (rs = .31-.71, all ps < .001; not shown in Table 2), indicating substantial co-occurrence and supporting their simultaneous entry in analyses.

**Table 2.**
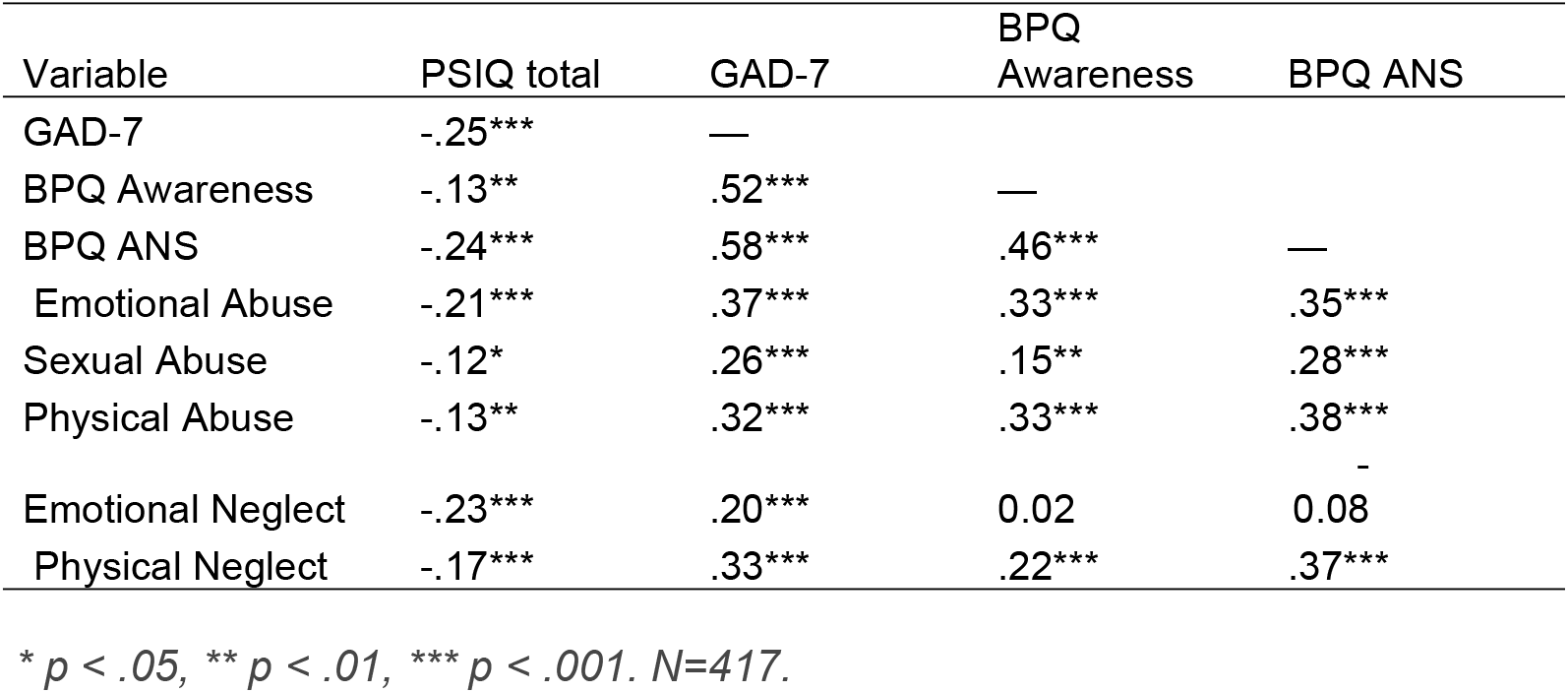
Correlations between childhood trauma subscales and mental imagery.

We also examined the interrelations among different types childhood trauma and mental imagery subscales. As shown in Supplementary Table 1, all PSIQ imagery subscales were strongly intercorrelated, and each showed consistent negative associations with CTQ trauma scores. In other words, across modalities (visual, auditory, smell, taste, touch, bodily, and feeling), higher trauma exposure was linked to lower imagery vividness. The uniformity of these effects suggests that imagery impairments in trauma are not confined to a single sensory domain but generalize across the imagery system.

### Hierarchical Multiple Regression

To identify which trauma types and symptom measures independently predict imagery vividness when accounting for shared variance, we conducted a hierarchical multiple regression analysis entering predictors in three sequential models (see Table 3).

**Table 3.** Hierarchical Multiple Regression Predicting Mental Imagery Vividness (PSIQ Total)

| Predictor | Model 1 |  |  | Model 2 |  |  | Model 3 |  |  |
| --- | --- | --- | --- | --- | --- | --- | --- | --- | --- |
|  | B | SE | p | B | SE | p | B | SE | P |
| **Demographics** |  |  |  |  |  |  |  |  |  |
| Age (years) | -0.005 | 0.048 | .922 | 0.005 | 0.048 | .917 | 0.005 | 0.048 | .916 |
| Gender (ref: Female) |  |  |  |  |  |  |  |  |  |
| Male | -0.510 | 1.568 | .745 | -0.648 | 1.550 | .676 | -0.375 | 1.509 | .804 |
| Non-binary | -21.416 | 14.664 | .145 | -21.781 | 14.499 | .134 | -17.925 | 14.103 | .204 |
| Other | -1.029 | 2.491 | .680 | -0.471 | 2.457 | .848 | -0.797 | 2.391 | .739 |
| **Childhood Trauma** |  |  |  |  |  |  |  |  |  |
| Emotional Abuse |  |  |  | -0.144 | 0.234 | .539 | -0.129 | 0.227 | .571 |
| Sexual Abuse |  |  |  | -0.073 | 0.172 | .672 | -0.017 | 0.168 | .920 |
| Physical Abuse |  |  |  | 0.262 | 0.252 | .299 | 0.382 | 0.245 | .120 |
| Emotional Neglect |  |  |  | -0.407 | 0.187 | .032* | -0.544 | 0.191 | .005** |
| Physical Neglect |  |  |  | 0.062 | 0.272 | .820 | 0.127 | 0.265 | .631 |
| **Interoceptive Symptoms** |  |  |  |  |  |  |  |  |  |
| BPQ-Body Awareness |  |  |  |  |  |  | -0.045 | 0.048 | .354 |
| BPQ-ANS |  |  |  |  |  |  | -0.254 | 0.087 | .004** |
| GAD-7 (Anxiety) |  |  |  |  |  |  | -0.336 | 0.172 | .052 |
| **Model Statistics** |  |  |  |  |  |  |  |  |  |
| R<sup>2</sup> | .006 |  |  | .065 |  |  | .126 |  |  |
| Adjusted R<sup>2</sup> | -.004 |  |  | .042 |  |  | .098 |  |  |
| $\Delta R^2$ | | | | .059 | | .001 | .061 | | <.001 |
| F | 0.58 |  | .677 | 2.83 |  | .003 | 4.48 |  | <.001 |

Note. $N = 410$ . $\Delta R^2$ and $p$ -values in Model Statistics row refer to the change from the previous model. \* $p < .05$ . \*\* $p < .01$ .

#### Model 1: Demographics Only

The first model included only demographic covariates (age, gender), accounting for 0.6% of variance in imagery vividness, F(4, 405) = 0.58, p = .677, Adjusted R^2^ = -.004. No demographic variables were significant predictors (all ps > .14).

#### Model 2: Addition of Childhood Trauma

In the second model, all five CTQ subscales measuring different types of childhood trauma were added. This significantly improved model fit, ΔR^2^ = .059, F(5, 400) = 5.08, p < .001, with the model now explaining 6.5% of variance, F(9, 400) = 2.83, p = .003,

Adjusted R^2^ = .042. Emotional neglect emerged as the only significant predictor (B = - 0.407, SE = 0.187, p = .032), while all other trauma types remained nonsignificant (all ps > .29).

#### Model 3: Addition of Interoceptive and Anxiety Symptoms

The final model added interoceptive symptom measures (BPQ Body Awareness, BPQ ANS) and current anxiety (GAD-7). This addition resulted in substantial improvement in model fit, ΔR^2^ = .061, F(3, 397) = 9.31, p < .001. The full model explained 12.6% of variance in imagery vividness, F(12, 397) = 4.48, p < .001, Adjusted R^2^ = .098.

Two predictors emerged as significant in the final model. Autonomic nervous system awareness (BPQ ANS) showed the strongest unique contribution (B = -0.254, SE = 0.087, t = -2.91, p = .004, 95% CI [-0.43, -0.08]), with emotional neglect remaining significant (B = -0.544, SE = 0.191, t = -2.85, p = .005, 95% CI [-0.92, -0.17]). Current anxiety showed a marginal effect (B = -0.336, SE = 0.172, t = -1.95, p = .052, 95% CI [-0.68, 0.00]). All other childhood trauma subscales (emotional abuse, sexual abuse, physical abuse, physical neglect) and BPQ-Body Awareness remained nonsignificant (all ps > .12). Diagnostic checks confirmed acceptable model assumptions. Multicollinearity was within acceptable limits (all VIFs < 1.8), the Durbin-Watson statistic (DW = 2.12) indicated no problematic autocorrelation, and residuals approximated normality despite minor deviations at the extremes (Shapiro-Wilk W = 0.952, p = .001), which are not problematic with N = 410.

The hierarchical regression revealed that when all predictors competed simultaneously (Model 3), autonomic nervous system awareness (BPQ ANS; B = - 0.254, p = .004) and emotional neglect (B = -0.544, p = .005) independently predicted imagery deficits. Most childhood trauma types showed significant bivariate correlations with imagery but became nonsignificant when entered with other predictors, suggesting their effects might operate through autonomic symptoms as a mediating mechanism. Importantly, non-significance in Model 2 does not indicate absence of effects, but rather suggests that these effects may be indirect. To formally test whether different types of childhood trauma impacts imagery through autonomic and anxiety pathways, we conducted parallel mediation analyses with all five trauma types as simultaneous predictors.

### Mediation Analysis

The parallel mediation model (N = 410, F(7, 402) = 7.67, p < .001; R^2^ = .118) simultaneously tested whether childhood trauma operates through autonomic symptoms, anxiety, or both pathways independently. Only Emotional neglect had a significant direct effect on PSIQ, while the others were fully mediated through BPQ ANS. Specifically, across all trauma types, BPQ ANS predicted reduced imagery (B =

-0.210, 95% CI [-0.35, -0.05], p = .004), while GAD showed only a marginal effect (B = -0.293, 95% CI [-0.60, 0.05], p = .076). Model diagnostics confirmed acceptable assumptions (VIF < 1.8, Durbin-Watson = 2.06-2.12). The significant mediation paths (with standardised beta coefficients) are shown in Figure 1.

**Figure 1.**
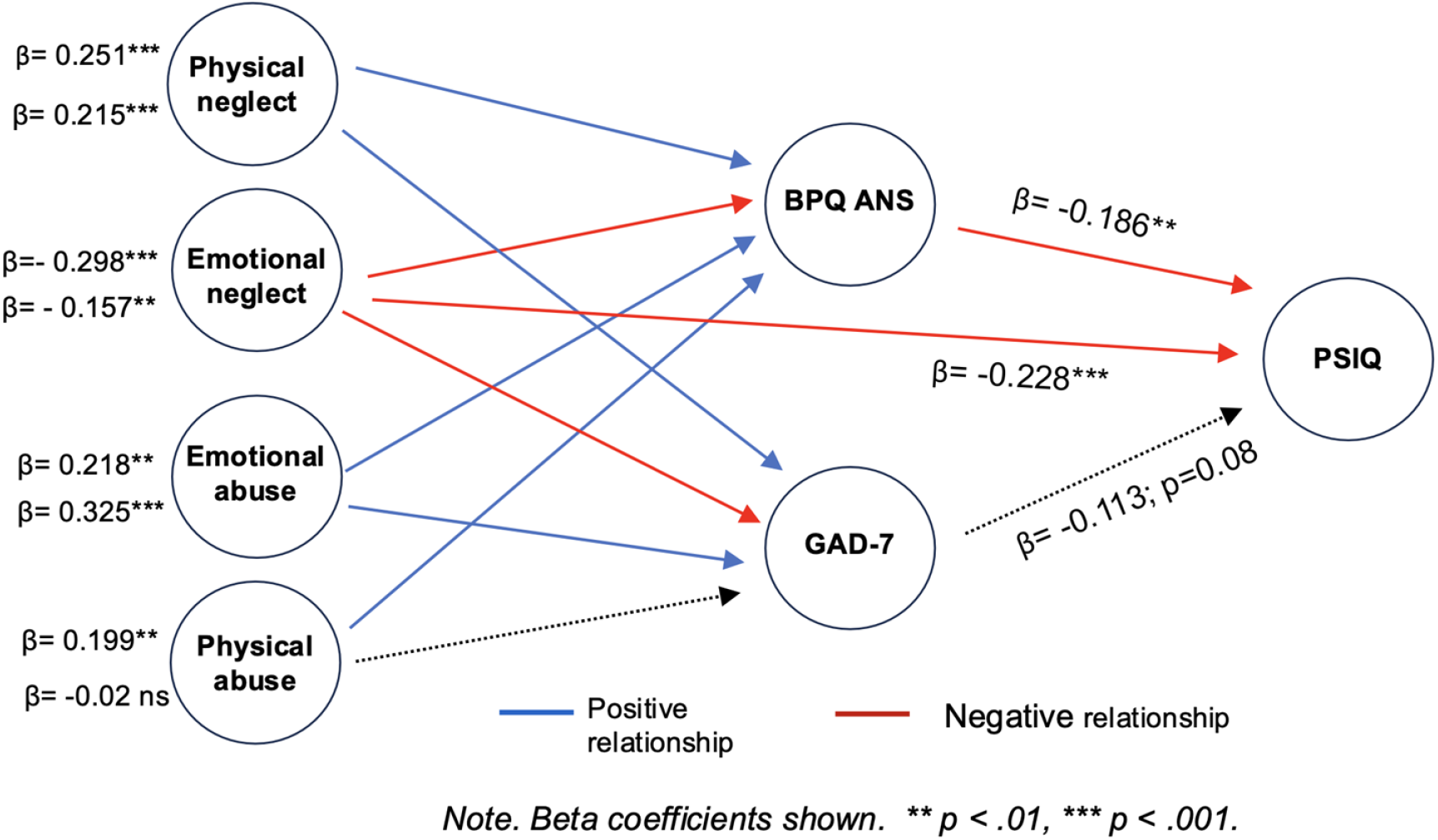
Parallel mediation model showing significant pathways from childhood trauma to mental imagery through autonomic symptoms (BPQ ANS) and anxiety (GAD. Values beside each predictor represent paths to BPQ ANS (top) and GAD-7 (bottom) Standardized coefficients (β) shown. * p < .05, ** p < .01, *** p < .001.

#### Indirect Effects Through BPQ ANS

Physical trauma types and emotional abuse predicted increased BPQ ANS scores (physical neglect: β = 0.251, p < .001; emotional abuse: β = 0.218, p = .002; physical abuse: β = 0.199, p = .004). For the GAD pathway, emotional abuse showed the strongest association (β = 0.325, p < .001), followed by physical neglect (β = 0.215, p< .001), emotional neglect (β = -0.157, p = .015), and sexual abuse (β = 0.108, p = .035). Physical abuse showed no significant association with GAD (β = -0.016, p = .818).

Four trauma types showed significant indirect effects through the BPQ ANS pathway: physical neglect (unstandardised B = -0.188, 95% CI [-0.38, -0.04], p = .019), physical abuse (B = -0.125, 95% CI [-0.31, -0.03], p = .042), emotional abuse (B = -0.117, 95% CI [-0.28, -0.02], p = .038), and emotional neglect (B = 0.147, 95% CI [0.04, 0.31], p = .014). Complete mediation was observed for emotional abuse and physical neglect, as their direct effects became non-significant when accounting for BPQ ANS (both ps .57). Sexual abuse showed no significant indirect effect through BPQ ANS (B = - 0.035, 95% CI [-0.12, 0.01], p = .276).

Emotional neglect demonstrated a contrasting pattern with a positive indirect effect alongside a strong negative direct effect (B = -0.607, p < .001; in the mediation model). This reflects emotional neglect’s unique association with decreased BPQ ANS (β = -0.298, p < .001), suggesting interoceptive blunting as a distinct mediating mechanism. The positive indirect effect indicates that lower autonomic awareness partially explains imagery deficits, while the substantial remaining direct effect suggests additional cognitive-representational mechanisms. Notably, this negative association with BPQ ANS emerged specifically in the mediation model when controlling for other trauma types (β = -0.298, p < .001), whereas the bivariate correlation was non-significant (r = -.08, p = .11). This suppression pattern indicates that emotional neglect’s unique effect of reducing autonomic awareness becomes apparent only after accounting for its co-occurrence with other trauma types that increase autonomic reactivity.

The GAD pathway showed no significant indirect effects for any trauma type (all ps .10), indicating that anxiety symptoms do not mediate the trauma-imagery relationship when controlling for autonomic symptoms.

## Discussion

The aim of this study was to examine the relationship between childhood trauma and mental imagery vividness. All trauma types were associated with reduced imagery across modalities, indicating domain-general impairments in mental simulation capacity. Emotional neglect and emotional abuse showed the strongest effects (rs = -.23 and -.21, respectively). Imagery was also negatively correlated with both autonomic symptoms (BPQ ANS; r = -.24) and body awareness (BPQ Body Awareness; r = -.13), with autonomic symptoms showing a substantially stronger association.

Trauma types showed distinct patterns of association with autonomic symptoms. Physical trauma (physical abuse, physical neglect) and emotional abuse showed robust positive correlations with the BPQ ANS subscale (rs = .35-.38), indicating heightened autonomic reactivity. Anxiety symptoms (GAD-7) were positively correlated with all trauma types.

Parallel mediation analyses revealed autonomic nervous system symptoms (BPQ ANS subscale) as the primary mediator linking childhood trauma to imagery deficits, while generalized anxiety (GAD-7) showed only borderline significant mediation. Various trauma types operated through the autonomic pathway, with physical neglect and emotional abuse showing complete mediation; their effects on imagery were entirely explained by heightened autonomic reactivity. Unlike other trauma types that predicted *increased* autonomic symptoms, emotional neglect predicted *decreased* autonomic awareness, reflecting interoceptive blunting. This contributed to imagery deficits through a positive indirect effect, but emotional neglect also retained a strong direct effect, indicating it impairs imagery through multiple mechanisms. Sexual abuse showed no significant indirect effect through autonomic symptoms, suggesting it may impact imagery through alternative pathways not captured by the current mediators.

### Mechanistic Pathways to Imagery Impairment

From a developmental perspective, childhood trauma involves repeated activation of threat-detection systems during critical periods of brain maturation. Chronic activation of the hypothalamic-pituitary-adrenal (HPA) axis and sympathetic nervous system during development can result in lasting autonomic dysregulation characterized by altered baseline arousal, heightened threat sensitivity, and impaired parasympathetic regulation (Dale et al., 2022; Dvir et al., 2014; Teicher & Samson, 2016). The present results suggest this autonomic dysregulation disrupts the interoceptive foundation required for vivid imagery.

In the interoceptive model of imagery, vivid imagery depends on the integration of sensory, affective, and interoceptive-autonomic inputs to construct coherent internal representations (Silvanto & Nagai, 2025). From this perspective, interoceptive signals provide the physiological scaffolding that grounds imagery in embodied experience, ensuring that internally generated representations remain vivid and somatically anchored rather than purely abstract. When interoceptive processing is dysregulated (whether through heightened or decreased autonomic reactivity) this integration is disrupted and imagery generation compromised. The domain-general nature of trauma-imagery associations across all sensory modalities (visual, auditory, olfactory, gustatory, tactile, bodily, emotional) is consistent with trauma affecting interoceptive systems that are suggested to provide a general foundation for imagery (Silvanto & Nagai, 2025).

Physical trauma types (physical abuse, physical neglect) were linked to heightened autonomic awareness, which in turn reduced imagery vividness through complete mediation. This pattern suggests that chronic interoceptive hypervigilance (i.e. the excessive monitoring of bodily sensations interpreted as potential threat) interferes with imagery through a two-stage process. First, trauma-related autonomic dysregulation produces heightened and often aversive bodily sensations. Second, when these internal signals become unpredictable, overwhelming, or persistently threatening, individuals may engage in compensatory detachment from interoceptive experience as a defensive strategy (Khalsa et al., 2018). This withdrawal from bodily awareness, observed in depersonalization and affective disorders (Sierra & Berrios, 2001; Zago et al., 2011), leads to imagery impairment.

Emotional neglect demonstrated a fundamentally different mechanism from other childhood adversities. Unlike physical abuse, emotional abuse, and physical neglect— which are typically event-linked and associated with autonomic hyperreactivity characteristic of acquired aphantasia (Gao et al., 2025)—emotional neglect operates as a chronic, pervasive environmental condition. Rather than triggering autonomic hyperreactivity, emotional neglect was associated with decreased autonomic awareness. Notably, this negative association with BPQ ANS emerged specifically in the mediation model when controlling for other trauma types (β = -0.298, p < .001), whereas the bivariate correlation was non-significant (r = -.08, p = .11). This suppression pattern indicates that emotional neglect’s unique effect of reducing autonomic awareness becomes apparent only after accounting for its co-occurrence with other trauma types that increase autonomic reactivity. This pattern showed a positive indirect effect on imagery deficits, such that lower autonomic awareness was associated with reduced imagery vividness.

This dissociation suggests distinct developmental pathways to imagery impairment. Event-linked adversities (physical abuse, emotional abuse, physical neglect) may trigger the hypervigilant, hyperreactive autonomic profile associated with acquired aphantasia—where traumatic experiences lead to defensive suppression of imagery accompanied by heightened somatic reactivity. In contrast, emotional neglect (characterized by persistent absence of emotional attunement and support for developing bodily awareness) may lead to developmental blunting of interoceptive processing. This reduced autonomic awareness pattern resembles the interoceptive profile observed in congenital aphantasia (Monzel et al., 2025), suggesting that chronic emotional deprivation during development may impair the maturation of interoceptive-imagery networks rather than triggering their defensive suppression.

However, emotional neglect also retained substantial direct effects independent of autonomic symptoms, indicating that interoceptive numbing explains only part of its impact. This dual-pathway pattern aligns with dimensional models distinguishing threat-based from deprivation - based adversity (McLaughlin et al., 2014). Deprivation of emotional responsiveness during development may additionally disrupt higher-order cognitive processes including self-referential processing (Philip et al., 2013; McLaughlin et al., 2015), and the formation of rich autobiographical representations (Fonagy et al., 2002). The combination of interoceptive and disrupted cognitive processes suggests emotional neglect impairs imagery through multiple, converging mechanisms.

Although both anxiety and autonomic symptoms were correlated with imagery deficits and childhood trauma, and despite substantial correlation between the two (r = .58, p< .001), only autonomic symptoms (BPQ ANS) significantly mediated the trauma-imagery relationship. GAD-7 showed only borderline significance in the mediator-to-outcome path and no significant indirect effects for any trauma type. This dissociation indicates that it is specifically the altered awareness of autonomic sensations that interferes with mental imagery, rather the cognitive rumination or worry components captured by GAD-7.

### Limitations

Several limitations should be noted. First, the cross-sectional design precludes causal inference. Although mediation analyses theoretically assume a causal direction, longitudinal studies are needed to confirm whether interoceptive dysregulation temporally precedes imagery deficits. Second, all measures relied on self-report, which may be influenced by factors such as memory bias, response style, and current psychological state. Objective indices of autonomic activity (e.g., heart rate variability) would provide stronger convergent evidence for the proposed autonomic pathway. Similarly, objective behavioral measures of imagery capacity (e.g., mental rotation tasks, imagery-based memory paradigms) would complement self-reported imagery vividness. Third, we did not assess other forms of psychopathology (e.g., depression, PTSD, dissociative symptoms) that may mediate or moderate trauma-imagery associations. For example, the effect of emotional neglect on imagery may at least partially operate through depressive symptoms or dissociative processes not captured by GAD-7.

## Conclusions

These findings demonstrate that childhood trauma impacts mental imagery through distinct autonomic mechanisms that vary by trauma type. Most physical trauma types and emotional abuse operated through heightened autonomic reactivity, with complete mediation observed for physical neglect and emotional abuse. In contrast, emotional neglect showed mediation through reduced autonomic awareness alongside the strongest direct effects on imagery, suggesting hyporeactive interoceptive blunting. These opposing patterns support the proposed dynamic range framework: both extremes of autonomic dysregulation (hyperreactive and hyporeactive) can disrupt the interoceptive signals necessary for vivid mental imagery. The findings reveal mechanistic heterogeneity in how different forms of early adversity impact cognition, with important implications for developing trauma-type-specific interventions targeting autonomic regulation.

